# Muscle nonshivering thermogenesis in a feral mammal

**DOI:** 10.1101/377051

**Authors:** Julia Nowack, Sebastian G. Vetter, Gabrielle Stalder, Johanna Painer, Maria Kral, Steve Smith, Minh Hien Le, Perica Jurcevic, Claudia Bieber, Walter Arnold, Thomas Ruf

## Abstract

While small mammals and neonates are able to maintain an optimal body temperature (T_b_) independent of ambient conditions by producing heat via nonshivering thermogenesis (NST) in the brown adipose tissue (BAT), larger mammals and other mammals lacking BAT were long believed to rely primarily on shivering and behavioural adaptations. However, recently, a second mechanism of NST was found in skeletal muscle that could play an important role in thermoregulation of such species. Muscle NST is independent of muscle contractions and produces heat based on the activity of an ATPase pump in the sarcoplasmic reticulum (SERCA1a) and controlled by the protein sarcolipin. To evaluate whether muscle NST could indeed play an important role in thermoregulation in species lacking BAT, we investigated the thermogenic capacities of new-born wild boar piglets. During cold exposure over the first 5 days of life, total heat production was improved while shivering intensity decreased, indicating an increasing contribution of NST. Sampling skeletal muscle tissue for analyses of SERCA activity as well as gene expression of SERCA1a and sarcolipin, we found an age-related increase in all three variables as well as in T_b_. Hence, the improved thermogenesis during the development of wild boars is not due to shivering but explained by the observed increase in SERCA activity. Our results suggest that muscle NST may be the primary mechanism of heat production during cold stress in large mammals lacking BAT, strengthening the hypothesis that muscle NST has likely played an important role in the evolution of endothermy.

## Introduction

The regulation of a high and stable body temperature (T_b_) independent of climatic conditions is one of the most important mechanisms that arose during the evolution of mammals and birds. After decades of intensive research, it is now well-understood how mammals possessing brown adipose tissue (BAT) - a specialised thermogenic organ - are able to maintain an optimal T_b_ even in cold environments by using nonshivering thermogenesis (NST) [reviewed in 1], However, only ~20% of endothermic birds and mammals actually possess BAT [2]. NST in BAT requires thermogenic functional uncoupling protein 1 (UCP1) that alters proton conductance in the inner mitochondrial membrane, leading to heat generation instead of ATP production [3, 4], Functional UCP1 has not been found, however, in marsupials or monotremes [5] and typically substantial amounts of BAT are present only in neonates of large-bodied mammals [6, 7, but see: 8]. Furthermore, a recent study has shown that UCP1-inactivating mutations have occurred in various mostly large-bodied placental mammals [9]. Large mammals, such as pigs, are likely able to cope well with cold exposure even without possessing BAT. Their juveniles, on the other hand, have high surface area-to-volume ratios and need a high capacity for heat production to maintain a constantly high T_b_ [10]. Inside the mother’s womb juveniles are protected against thermal variations and they are exposed to a cold environment for the first time after birth. In fact, it is known that neonates of many large-bodied species in which BAT depots are negligible in adults, possess large amounts of BAT [6, 7].

It was long believed that species without thermogenic functional UCP1 rely solely on shivering, a process that on its own is insufficient for the maintenance of a stable T_b_ during cold exposure in UCP1 knockout mice [11]. However, a second mechanism of NST in muscle, which had been studied in vitro for a long time [12-15], has recently been shown to play an important role in thermoregulation in mice lacking functional BAT [11], The mechanism is so far only confirmed as an additional form of NST in rodents in which the wildtype possesses BAT [11, 16-20], but assumed to occur in all mammals [2, 21]. Furthermore, muscle NST is discussed as a heat production mechanism that played an important role for the evolution of a high T_b_, i.e. endothermy, in mammals [2, 21], as endothermy in this group evolved before UCP1-mediated NST [22].

In short, muscle NST is based on the activity of the Ca^2+^-ATPase pump in the sarcoplasmic reticulum (SERCA). SERCA1a, the major isoform occurring in skeletal muscle [23], is involved in muscle contraction via the transport of Ca^2+^-ions from the cell lumen into the sarcoplasmic reticulum [24, 25]. But ATP hydrolysis by SERCA1a can be uncoupled from actual transmembrane transport of Ca^2+^ by the regulatory protein sarcolipin (SLN) causing the release of the Ca^2+^-ions bound to SERCA back to the cytoplasmic side of the membrane (so called “slippage”) rather than into the sarcoplasmic reticulum [reviewed in 26]. This results in increased ATP hydrolysis and heat production in muscle through SERCA1a activity without actual Ca^2+^-transport and without muscle contraction [13, 26-29]. Studies on laboratory mice have shown that muscle NST can compensate for the loss of UCP1, but muscle NST in wildtype mice is largely masked by heat production in BAT [30, 31]. This raises the question whether the thermogenic capacity of muscle NST alone can enable mammals to maintain a stable T_b_ under cold conditions.

We investigated whether muscle NST is an important and effective mechanism to generate heat in wild-type mammals lacking BAT, using new-born wild boars *(Sus scrofa),* naturally lacking BAT [32] and the UCP1-dependent NST [33, 34] as model species. Importantly, juvenile wild boars are born in early spring, when ambient temperatures (T_a_) can still be around or below 0°C and thermoregulatory demands are high. We firstly hypothesized that, if NST plays a significant role for thermogenesis in piglets, heat production should increase during cold exposure, whereas shivering should remain constant or may even decrease. We secondly hypothesized that, if heat production is based on muscle NST, expression levels of SERCA1a and sarcolipin as well as SERCA activity should be elevated during cold exposure.

## Material and Methods

### Experimental setup

Piglets were born in March 2017 by five sows kept and bred in outdoor enclosures at the Research Institute of Wildlife Ecology (48.22° N, 16.28° E) of the University of Veterinary Medicine Vienna in Austria. Each sow was provided with a roofed shelter outfitted with straw in which they gave birth. Shelters were equipped with IP-cameras and activity of the sows was monitored to know the exact time of birthing. All animals were exposed to natural T_a_ but piglets typically huddled with each other or with adults. Mean daily T_a_ during the 5 days of our study ranged from 5.9 °C to 14.7 °C and nightly minimum T_a_s were between 1.5 °C and 9.4 °C.

We temporarily removed 19 of 29 new-born wild boars from their mothers within the first 24 hours after birth (age 8.5-24 h) and again four days later. Rectal T_b_ was taken within 5-10 minutes after removal from the mother by inserting a thermometer with rectal gel approximately 2 cm into the rectum. Piglets were weighed to an accuracy of 1 g (Sartorius, Göttingen, Germany) and equipped with a custom-made acceleration logger (see below) that was firmly attached to their abdomen with a cohesive bandage (Henry Schein, New York, USA) before being placed individually into metabolic chambers (40 x 25x 22 cm, volume: 20 l). The metabolic chambers were located inside a walk-in climate chamber that was set to + 10 °C (mean 10.3 ± 0.7 °C). Individuals were exposed to cold for approximately 60-90 min and heat production and shivering intensity during cold exposure were determined (see below). After measurements rectal T_b_ was determined again and piglets were transferred to a surgery room where muscle biopsies were taken (see below).

### Metabolic measurements

All experiments were conducted between 0800h and 1630h. Metabolic measurements lasted for 60-90 min (depending on the activity level of the animal). Animals were allowed to acclimate to cold conditions for 10 min, before the measurement was started. Energy expenditure was determined by measuring the rate of O_2_ consumption (VO_2_) as a proxy of metabolic rate using an O_2_ and CO_2_ analyser (Servopro 4100, Servomex, Crowborough, UK). The metabolic chamber was connected to the analyser (pull mode; order: metabolic chamber, pump, needle valve, flow meter, O_2_ analyser, CO_2_ analyser) with airtight tubes. Water vapour was removed from the air prior to analysis using silicagel. A gas switch allowed measurement of air from six metabolic cages and one reference air channel for one minute each. The analyser was calibrated once a week using a high precision gas-proportioning pump (H. Wösthoff, Bochum, Germany, type 55A27/7a). Air was continuously drawn through the cages with pumps at a flow rate of 250 on day 1 and - 330 l h^-1^ on day 5. Flow rate through each metabolic chamber was measured using calibrated thermal mass-flow meters (FMA 3100, Omega Engineering, Stamford, CT, USA). VO_2_ was calculated by a self-written R program [35] using equation 10.6 by Lighton [36] and converted to heat production (watts) assuming 20.1 J ml^-1^ O_2_ consumption [37], VO_2_ were computed from the mean of the three lowest consecutive values per measurement. To standardize measurements we only used the first 30 min of the measurement for calculations.

### Shivering

We successfully measured shivering on day 1 and day 5 of 12 cold-exposed piglets by attaching a custom-made acceleration logger (three axis acceleration sensor ADXL345; 3 x 1 x 45 mm; Li-polymer rechargeable Battery LP-402025-1S-3; weight with battery of 6.4 g). The sensor of our acceleration logger measured acceleration in three orthogonal axes (x, y, and z). Sampling rate of the acceleration sensor was set to 1600 Hz, acceleration range was ± 16 *g* and had a resolution of 4 mg per axis. The battery had a capacity of 155 mAh limiting the runtime to approx. 6 hours, which was sufficient for our needs. We analyzed the acceleration sensor data entirely in the frequency domain. For this purpose, we calculated the resulting acceleration from the x, y and z-components:

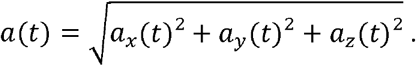

Then we subtracted the mean acceleration stemming from gravitation and applied a Fast Fourier Transformation (FFT) in R (V3.4.2) [38] *(fft* in package ‘signal’ [39]) to consecutive segments of 6 second intervals in the time domain (see supplementary figure S1). From every FFT output we extracted the following parameters:

1. Maximum Amplitude (m/s^2^)
2. Frequency at Maximum Amplitude (in Hz)

Exposure to 10°C without the opportunity to huddle with conspecifics caused clearly visible shivering with short intermitted phases of suspension of shivering. To calculate shivering intensity (the maximum amplitude) we used the same time frame as for metabolic rate calculations, i.e. a total of 30 min, starting after 10 min of acclimatization. The frequency of muscle activation during shivering thermogenesis is correlated with body mass in mammals [40] and is reported to be: log*f*= 1.85 – 0.18 logm, where *f* is the mean shivering frequency in Hz and *m* is the mean body mass in g [40]. Based on this equation the shivering frequency of day 1 to day 5 old piglets with a mean body mass of 1700 g ± 306 g is predicted to be 18.55 Hz. We therefore restricted our analyses to frequencies between 10 Hz and 30 Hz and excluded frequencies above or below this range from the analysis. Measured mean shivering frequency at maximum amplitude for piglets was with 17.97 ± 1.66 Hz (N= 24) in the predicted range and did not differ between days (χ^2^= 0.540, df= 1, p= 0.462).

### Biopsies

80-100 mg muscle tissue was taken with a biopsy needle from the thigh region of the piglets *(Musculus semimembranosus)* under standard surgical conditions. The procedure was conducted under general anaesthesia in combination with local anaesthesia. The anaesthesia was induced by placing a mask over the piglets mouth and nose, using an Isofluran gas (Isofluran, Isoba, MSD animal health, Vienna, Austria) inhalation machine and medical oxygen. Pain relief was achieved by injecting local anaesthetics (Lidocain, Xylocain 2 % with Epinephrin 1:100,000. Mibe GmbH, Brehna, Germany) 10 min prior biopsy in the proximal region of the *M. semimembranosus* and the skin. Anti-inflammatory and pain relief treatment was furthermore gained by injecting Meloxicam 10 min prior biopsy intramuscularly (Meloxicam 0.4 mg/kg, Metacam 2% inj., Boehringer Ingelheim Vetmedica, Ingelheim, Germany). During the entire procedure, vital parameters (respiration rate, peripheral haemoglobin oxygen saturation as measured by pulse oximetry (SpO2), heart rate, T_b_) and anaesthetic depth were monitored. The skin and muscle fascia incisions were closed separately in two layers with absorbable sutures (Surgicryl USP 2/0 PGA, SMI AG, Hünningen, Belgium). Piglets were marked with an ear tag for further individual recognition. After the biopsy the juveniles were monitored and kept in a warm environment for recovery from anaesthesia until they were moved to their enclosure. Piglets were fed with 1 mL of glucose solution orally (50 % glucose infusion, B.Braun AG, Melsungen, Germany) before being returned to their mothers. Piglets were never kept away from their mother for more than 3-4 hours. The muscle tissue sample was split for biochemical as well as genetic analyses. We were able to obtain samples on both days of 17 piglets. The two animals for which we only obtained one sample were excluded from further analyses regarding SERCA1a and SLN. For two further animals tissue material was too small to be used for biochemical analyses, which reduced the sample size for SERCA activity to N= 15.

### Biochemical analysis

Approximately 50 mg of muscle tissue of 15 piglets were snap frozen within 10 min and later stored at -80°C until used to prepare muscle homogenates according to the procedures described in Giroud et al. [41]. Homogenates were used to measure SERCA activities by a standard coupled enzyme assay, in which the rate of SERCA ATP hydrolysis was calculated from spectrophotometric recording (method previously described by Simonides et al. [42]). While this assay is measuring overall ATPase activity of all isoforms of SERCA present in the sample, SERCA activity is assumed to primarily reflect SERCA1a activity, because SERCA1a is the major isoform found in fast twitching muscles, such as *M. semimembranosus* [23]. SERCA activity was divided by total protein concentration (determined with Bradford method [43]) in order to normalize the ATPase activity values for variations in total protein concentrations among samples. More details on isolation of muscle homogenates and SERCA activity measurements can be found in the supplementary methods.

### Genetical analyses

Further, 30 mg muscle tissue of 17 piglets were directly stored in RNAlater^®^, kept in the fridge for 24 hours and stored at -80°C until RNA was extracted using the RNeasy Fibrous Tissue Mini Kit (Qiagen, Hilden, Germany). Gene expression levels for SLN and SERCA1a were analysed from cDNA via Droplet Digital PCR (ddPCR™). RNA was reverse-transcribed with MultiScribe™ Reverse Transcriptase (High Capacity cDNA Reverse Transcription Kits, ThermoFisher Scientific) using random hexamer primers. Primer sequences for the target gene, SLN, were available from Vangheluwe et al. [44], No primer sequence was available for SERCA1a so we designed suitable primers from the reference sequence NM_001204393.1 with the assistance of the NCBI primer design tool [45]. Primers for candidate reference genes *Hypoxanthirie Phosphoríbosyltransferase 1 (HPRT1)* and *Glucuronidase Beta (GUSB)* were available from [46]. All primer sequences as well as additional methodological details can be found in the supplementary methods and table S1. Data acquisition was accomplished by the QX200™ Droplet Reader (Bio-Rad), and analysed using the Bio-Rad Droplet Digital™ PCR QuantaSoft software. Expression levels are given as the relative ratio of the concentration (copies/μl) of the assay target gene over the concentration of the reference gene. Importantly, SLN protein expression has been shown to correlate well with SLN mRNA expression [31].

### Data analyses

Data are presented as mean ± 1 S.E.; N denotes the number of individuals. Data were analysed in R (V3.4.2)[38]. We first tested whether SERCA1a gene expression, SLN gene expression, SERCA activity and T_b_ increased with age (using group “day 1” and “day 5”) by employing a linear-mixed effect model followed by type II sum-of-squares ANOVA (lme in library ‘nlme’ [47]; Anova in library ‘car’ [48]). All models were corrected for non-independence by including the individual’s mother as well as the individual’s ID as nested random effects. To investigate whether SERCA activity was explained by SLN and SERCA1a gene expression, we additionally computed linear-mixed effect models including random effects as described above (pseudo r^2^ was calculated using sem.model.fits in library ‘piecewiseSEM’ [49]). For the cold exposure experiments we tested whether there was an age depended change in heat production, shivering intensity and shivering frequency as described above. In tests for changes of total heat production, we included body mass as a covariate. Mass-specific rates of heat production were computed for graphical presentation, but not used for statistical tests. To test T_b_ regulation over the time of cold exposure we computed a linear mixed effect model, again with ID and mother as random effects, T_b_ at the end of exposure as the response variable, and day (1 or 5), initial T_b_, and duration of cold exposure (60-90 min) as fixed effects. We used Shapiro-Wilk tests to access the normality of model residuals. If needed, data were Box-Cox transformed.

## Results

### Cold exposure experiment

Heat production during cold exposure did not significantly differ between day 1 and day 5 when adjusted for the increase in body mass (χ^2^= 1.92, df= 1, p= 0.165; N= 19; Fig. 3a), although shivering intensity decreased between both measurements by 50% (χ^2^= 9.00, df= 1, p= 0.003; N= 12; Fig. 3b). T_b_ after cold exposure was dependent on T_b_ at start of the experiment (Tab. 2; χ^2^= 8.01, df= 1, p=0.0046), but did not significantly differ between the days (χ^2^= 3.33, df= 1, p= 0.0679) and was not dependent on the duration of cold exposure (χ^2^= 1.86, df= 1, p= 0.173).

**Table 1:** Comparison of physiological SERCA activity and the amount of mRNA expression of SLN and SERCA1 at day 1 and day 5 after birth. Sample sizes of tested wild boar piglets are given in brackets. Different letters indicate significant differences. Statistical test results are reported in the text. * Gene expression is reported in copies per μl target gene/copies per μl reference gene (HPRT1). ^+^ SERCA activity is reported as ATP hydrolyses per minute and mg total protein.

|  | Body mass (g) | SERCA activity<sup>+</sup> | SERCA1a mRNA expression* | SLN mRNA expression* |
| --- | --- | --- | --- | --- |
| Day 1 | 1411.9 ± 16.5 (N=19) | 0.36 ± 0.07 (N=15)<sup>a</sup> | 109.6 ± 19.7 (N=17)<sup>a</sup> | 40.9 ± 7.8 (N=17)<sup>a</sup> |
| Day 5 | 1987.6 ± 29.0 (N=19) | 0.96 ± 0.12 (N=15)<sup>b</sup> | 320.1 ± 47.2 (N=17)<sup>b</sup> | 93.7 ± 14.5 (N=17)<sup>b</sup> |

**Table 2:** Body temperature (T_b_) regulation during cold exposure at 15°C. (mean ±SD). Statistical test results are reported in the text.

|  | Day 1 | Day 5 |
| --- | --- | --- |
| $T_b$ (°C) before cold exposure | 38.6 ± 0.09 (N=19) | 39.2 ± 0.07 (N=19) |
| $T_b$ (°C) after cold exposure | 38.6 ± 0.09 (N=19) | 39.0 ± 0.07 (N=18) |

**Figure 1:**
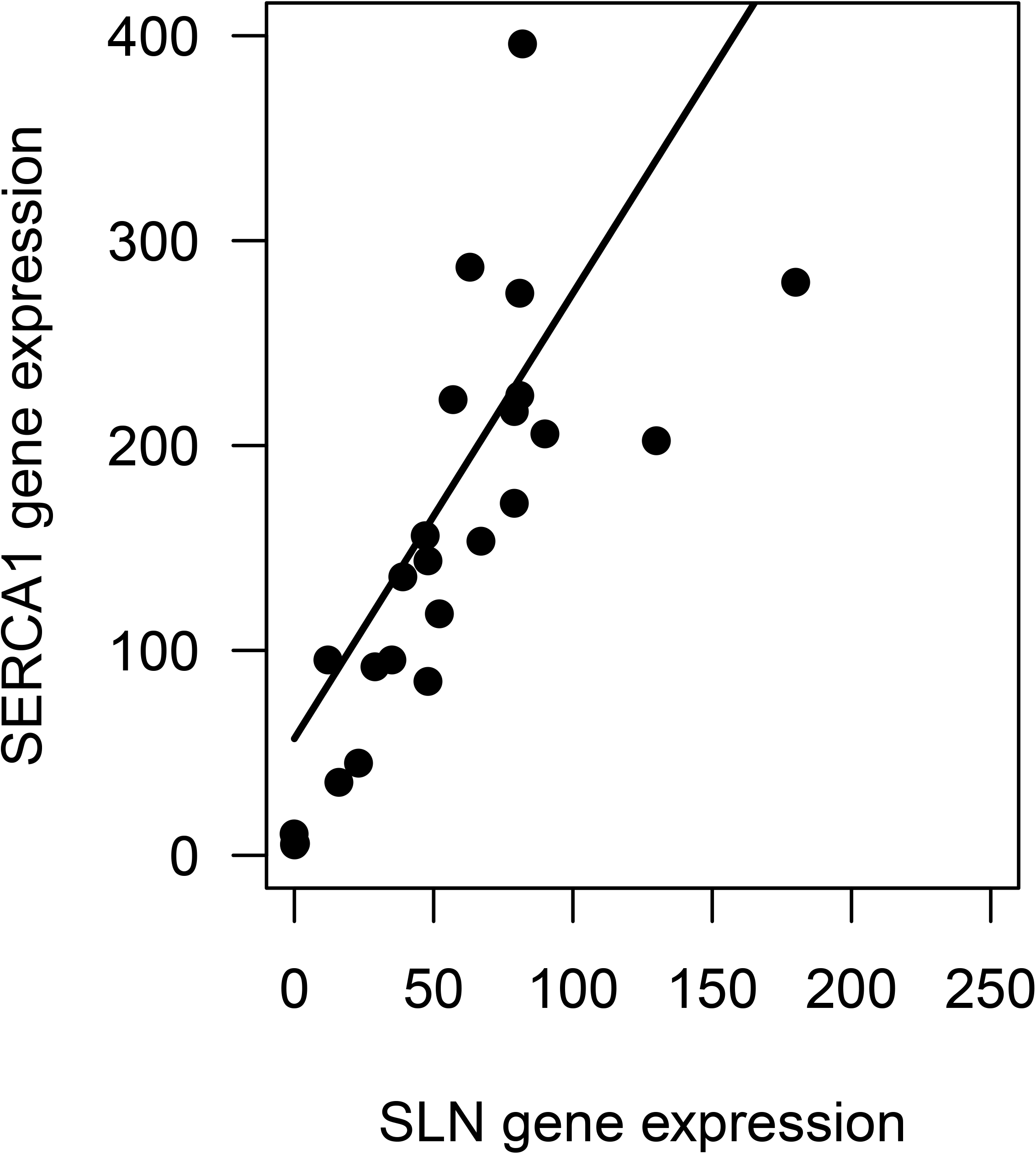
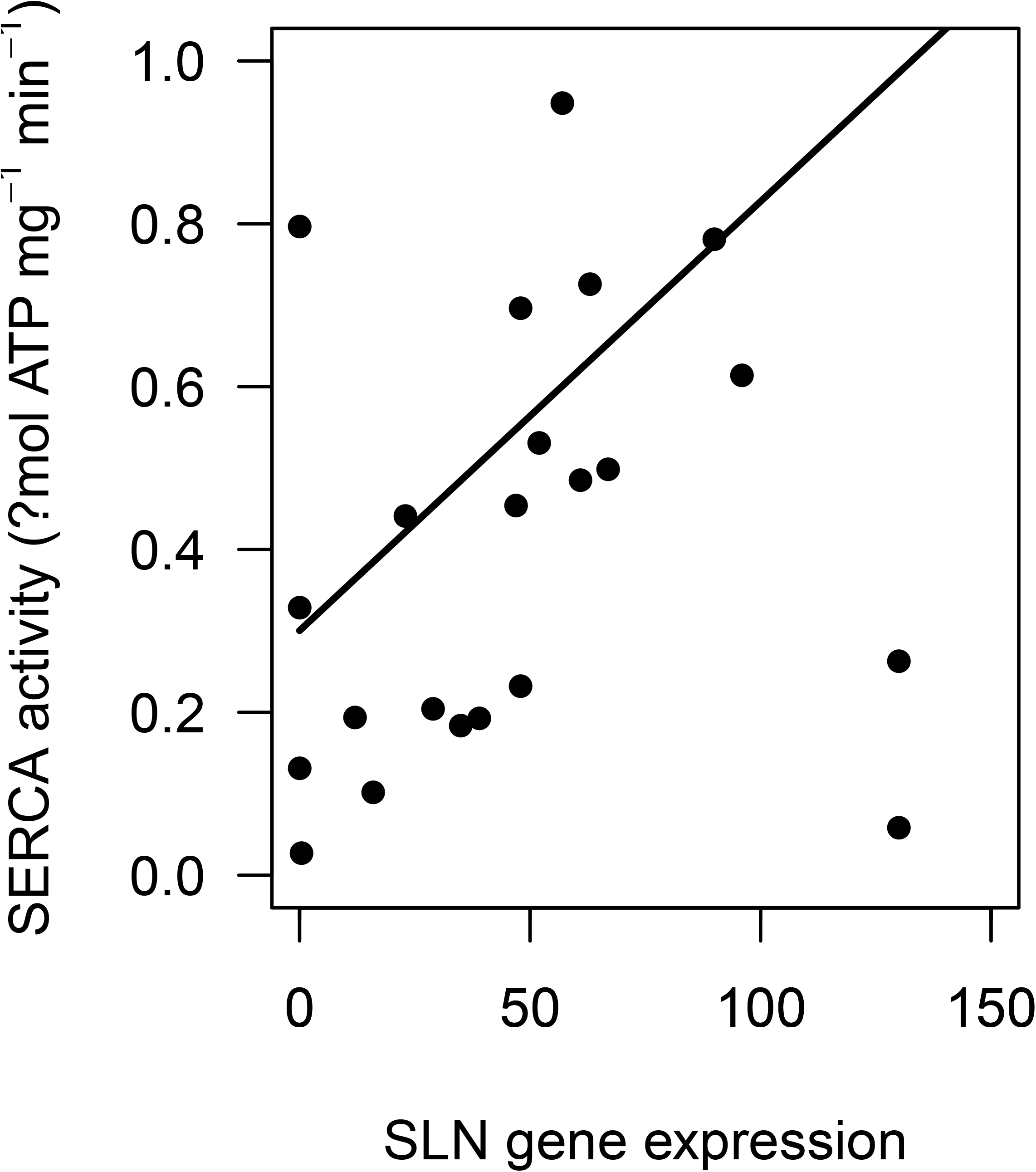
Correlation of SERCA1a and SLN gene expression. The expression of both genes was significantly correlated (χ^2^=32.2, df=l, p<0.0001, marginal pseudo r^2^: 0.51). Gene expression is reported in copies per μl target gene/copies per μl reference gene (HPRT1).

**Figure 2ab:**
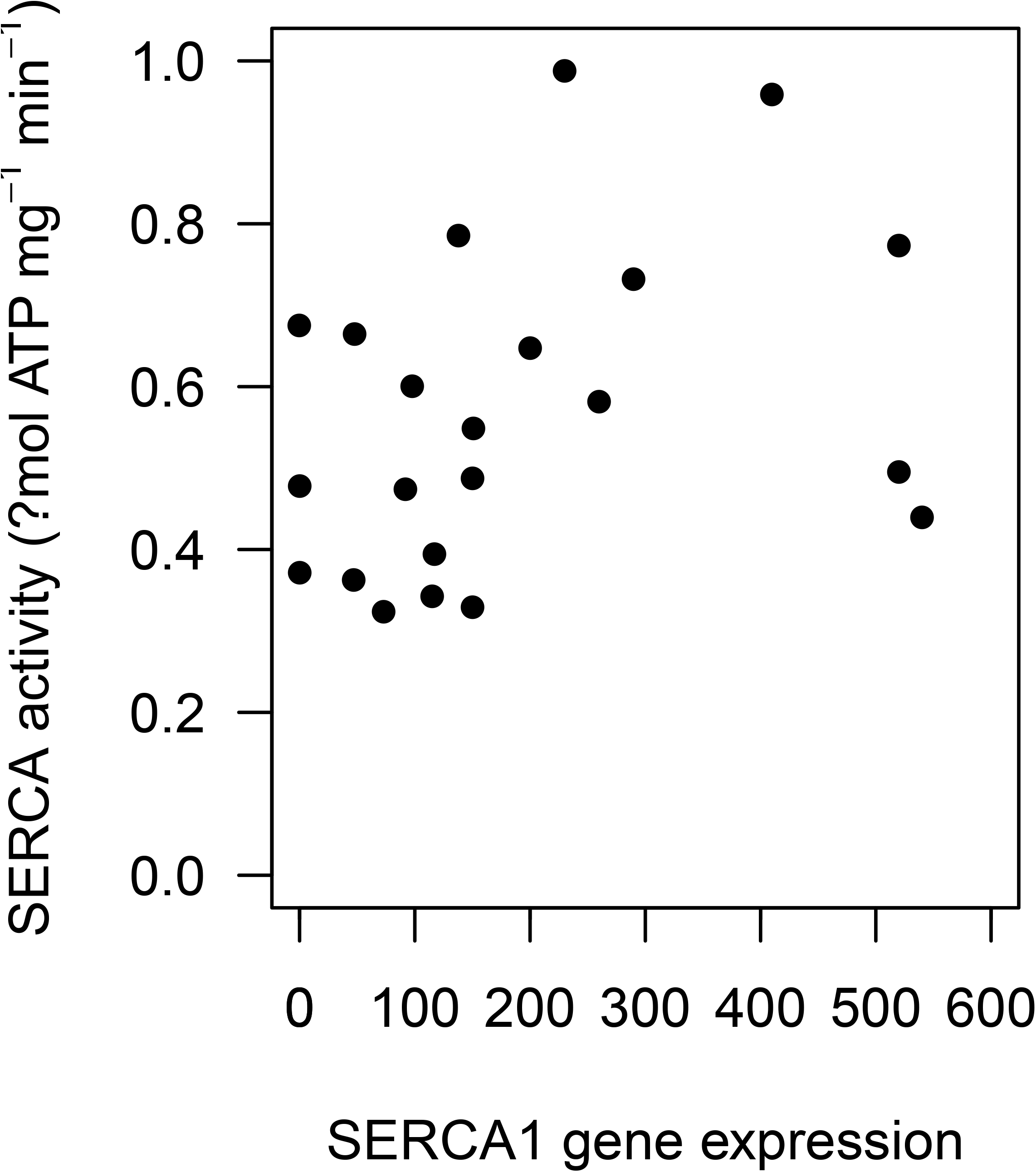
Partial regression plots of the effect of SLN and SERCA1a gene expression on SERCA activity. The increase of SERCA activity was linked to SLN gene expression (χ^2^=6.57, df=1, p=0.01), while SERCA1a gene expression had no significant effect on SERCA activity when tested together with SLN (χ2=0.002, df=1, p=0.967). Gene expression is reported in copies per μ1 target gene/copies per μl reference gene (HPRT1), SERCA activity as ATP hydrolyses per minute and mg total protein.

**Figure 3ab:**
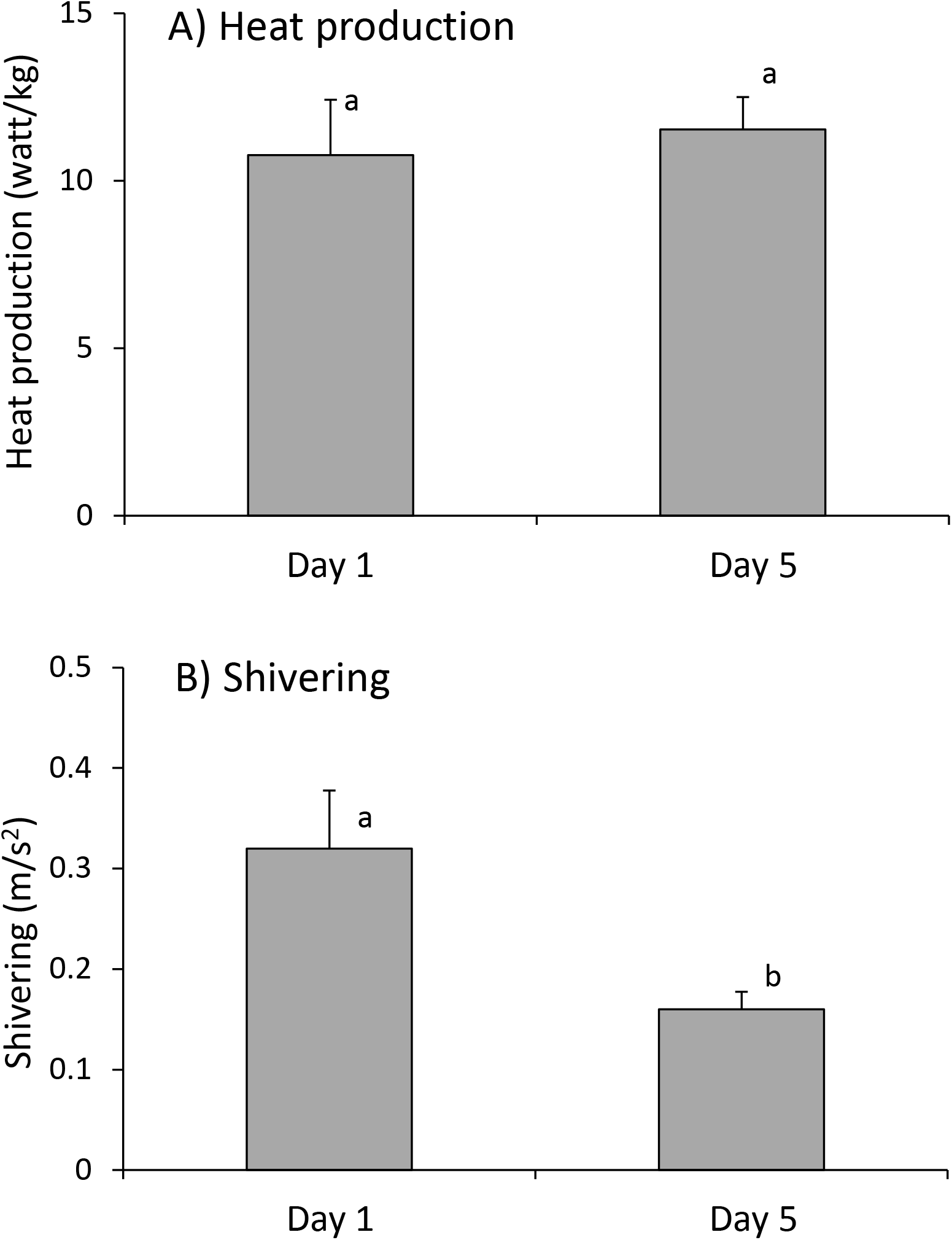
Change in heat production and shivering between day 1 and day 2. Different letters indicate significant differences. Heat production, adjusted for the increase in body mass, did not significantly differ between days (N=19; χ^2^=1.92, df=1, p= 0.165), whereas shivering significantly decreased (N=12; χ^2^=9.00, df=1, p=0.003).

### SERCA1a, SLN and SERCA activity

SERCA activity, as well as SERCA1a and SLN mRNA expressions increased significantly from day 1 to day 5 of the juveniles’ life (Tab. 1; SERCA1: χ^2^= 15.29, df= 1, p<0.0001; SLN: χ^2^= 9.63, df= 1, p= 0.002; SERCA activity: χ^2^= 16.81, df= 1, p<0.0001) and SERCA1a and SLN expression levels were positively correlated (χ^2^= 32.2, df= 1, p<0.0001, marginal pseudo r^2^: 0.51; Fig.1). Importantly, the age-related increase of SERCA activity was linked to the increase in SLN gene expression, when tested in a single-predictor model (χ^2^= 13.68, df= 1, p<0.001) as well as in multiple regression together with SERCA1 (χ^2^= 6.57, df= 1, p= 0.01; Fig.2a); SERCA1a gene expression was only significantly influencing SERCA activity when used in a single-predictor model (χ^2^= 7.23, df= 1, p= 0.007), but not when tested together with SLN (χ^2^= 0.002, df= 1, p= 0.967; Fig. 2b). T_b_ of piglets increased significantly from day 1 to day 5 (χ^2^= 41.01, df= 1, p<0.001; Tab.2) and this increase was linked to SERCA activity (χ^2^= 5.22, df= 1, p= 0.02).

## Discussion

Our study revealed that shivering intensity decreases from day 1 to day 5 in juveniles exposed to 10°C, while heat production during cold is increasing proportional to body mass and the level at which T_b_is maintained increases. This result is clear evidence for an increasing contribution of NST to thermogenesis during cold exposure in piglets over the first days of life. The finding that simultaneously, SERCA activity and the expression of SERCA1a and SLN were recruited points to muscle NST and increased SERCA activity as the principal source of heat production, as we can rule out UCP1-mediated NST in this species.

Our statistical analysis suggests that the increase in SERCA activity between day 1 and day 5 was mainly due to an SLN controlled up-regulation of ATP hydrolysis by SERCA1a instead of an increase in SERCA1a molecules. This is in accordance with previous data on mice in which upregulated SLN expression led to an increasing contribution of SERCA-based Ca^2+^-slippage to heat production [11, 17].

In mice and rats, in which neonates are born blind and naked (‘altricial’ neonates), the transcription and translation of SLN is highest after birth and gradually decreases with development when kept at normal housing conditions (~23°C) [16, 31], while the amount of BAT is successively recruited [1]. However, when kept under cold conditions juvenile mice keep SLN up-regulated for improved thermoregulatory capacity [31], suggesting that both mechanisms of NST are necessary for an effective maintenance of a high T_b_ during cold exposure in new born rodents. In adult mice, however, both mechanisms of NST, UCP1-mediated as well as muscle NST, can compensate for the loss of one system [50]. In contrast, our data indicate that muscle NST, in combination with some shivering, is already sufficient to maintain a stable T_b_ for short-term cold exposure in juvenile wild boar, which are markedly larger than juvenile mice and are already born with fur and much better thermogenic abilities (‘precocial’ juveniles). Interestingly, in other precocial species, such as sheep and goats that possess functional UCP1, BAT is recruited already before birth [1], Furthermore, reconstituted function of UCP1 can further improve thermoregulatory function of cold exposed 6-months old Bama pigs, a cold-sensitive pig breed.

Our finding that muscle NST is involved in thermoregulation of juvenile wild boars and allows a near stable T_b_ even during short-term cold exposure supports the hypothesis that muscle NST may be the primary mechanism of heat production during cold-exposure in large mammals lacking BAT. While the evolution of BAT has often been related to the ability of small placental mammals to colonize colder habitats [4, 51, 52], a recent study has shown that UCP1-inactivating mutations have occurred in at least eight of the 18 placental mammalian orders, mainly in larger-bodied species [53], such as pigs. It therefore appears that the combination of shivering and muscle NST is sufficient for heat production in large mammals. Pigs, for example, likely lost UCP1 function and the ability to use BAT for thermoregulation because of absent or only weak selection for this mechanism in a warm climate [54]; all species except the wild boar live only in tropical or subtropical habitats. In addition to heat production via muscle NST, wild boar apparently evolved compensatory mechanisms to cope with adverse thermal conditions in northern habitats, such as larger adult body size [55], building insulating nests for offspring, and synchronizing reproduction within social groups or enabling piglets to huddle in large groups of combined litters [54, 56]. Behavioural thermoregulation is less energetically costly than NST [57] and a study on winter mortality of juvenile wild boar has shown that the negative effects of cold winters can be compensated by high availability of food resources [55].

In addition to our finding that SLN-mediated NST in skeletal muscle is involved in piglet thermoregulation, recent studies on domestic pig breeds suggest that SERCA2b (another isoform of SERCA) and UCP3 might also influence pig thermoregulation [58, 59]. However, so far the importance of both mechanisms is unclear [e.g. 2] and the evolution of a compensatory mechanism after the pigs colonized cold habitats is likely [58], while muscle NST is discussed as a potentially evolutionary old heat production mechanism [e.g. 2, 21]. Whether and to what extend domestic pig breeds also possess muscle NST remains speculative. While piglets of wild boar are accustomed to deal with temperatures around or below zero degrees, domestic pigs are kept under warm conditions (20-35°C). Therefore, it cannot be ruled out that the extreme susceptibility of pigs to cold is partly due to inadvertent selection against high thermogenic capacity during domestication. Previous studies on thermoregulation of juvenile domestic pigs have also found that shivering intensity decreased during the first days after birth while heat production and blood flow to muscles simultaneously increased [60-62], While this was originally attributed to an increase in shivering efficiency [60, 61], it seems questionable whether an increased thermogenesis by increased efficiency of shivering is physically possible. Our data now suggest that the improved thermogenesis found in domestic pigs, similarly to wild boar, was not due to an increase in shivering efficiency, but explained by an increase in muscle NST.

Taken together, our data show for the first time that muscle-based NST via SERCA1a plays a role in the thermoregulation of wild type mammals lacking BAT and that muscle NST can replace UCP1-mediated NST. The function of UCP1 as a thermogenic protein has occurred after the divergence between placental and marsupial mammals [22], suggesting that the evolution of endothermy in ancestral mammals was independent of heat production in BAT. Although the earth was likely warmer, ancestral mammals still would have experienced daily and yearly fluctuations in Ta, likely similar to temperatures found in tropical areas today, which can get rather cold during the night. Therefore, while heat produced as a by-product of metabolic processes as well as basking [63] would have allowed the establishment of a stable T_b_ during a big part of the day, muscle NST was likely important during the colder night hours.

### Ethics statement

The study was approved by the institutional ethics and animal welfare committee and the national authority according to §§ 26ff. of Animal Experiments Act, Tierversuchsgesetz 2012 – TVG 2012 (BMWFW-68.205/0171-WF/V/3b/2016).

### Data accessibility

The data will be made available at figshare upon acceptance of the publication.

## Acknowledgements

We thank Peter Steiger and Michaela Salaba for their help with animal maintenance, Joy Einwaller, Jessica S. Cornils and Arne Müller for help during the experiments, Omid Hekmat and Michael Hämmerle for biochemical analyses and Martin Olesch, Thomas Paumann and Radovan Kovacki for infrastructural support. The study was supported by funding from the Humboldt foundation to JN and by the University of Veterinary Medicine, the City of Vienna, and the Government of Lower Austria.

## Author Contributions

JN and TR designed the experiments, JN conducted the experiments, analysed the data and wrote the manuscript, SV was involved in the performance of the experiments and helped with statistical analyses, GS and JP performed the biopsies, ML and OH conducted the biochemical analyses, MK and SS conducted the genetic analyses, JP designed the accelerometers and computed the shivering intensities, CB designed the enclosures and organised logistics, CB and WA were involved in the discussion of the experimental plan. All authors commented on the manuscript.

